# The influence of cortical depth on neuronal responses in mouse visual cortex

**DOI:** 10.1101/241661

**Authors:** Philip O’Herron, John Woodward, Prakash Kara

## Abstract

With the advent of two-photon imaging as a tool for systems neuroscience, the mouse has become a preeminent model system for studying sensory processing. One notable difference that has been found however, between mice and traditional model species like cats and primates is the responsiveness of the cortex. In the primary visual cortex of cats and primates, nearly all neurons respond to simple visual stimuli like drifting gratings. In contrast, imaging studies in mice consistently find that only around half of the neurons respond to such stimuli. Here we show that visual responsiveness is strongly dependent on the cortical depth of neurons. Moving from superficial layer 2 down to layer 4, the percentage of responsive neurons increases dramatically, ultimately reaching levels similar to what is seen in other species. Over this span of cortical depth, neuronal response amplitude also increases and orientation selectivity moderately decreases. These depth dependent response properties may be explained by the distribution of thalamic inputs in mouse V1. Unlike in cats and primates where thalamic projections to the granular layer are constrained to layer 4, in mice they spread up into layer 2/3, qualitatively matching the distribution of response properties we see. These results show that the analysis of neural response properties must take into consideration not only the overall cortical lamina boundaries but also the depth of recorded neurons within each cortical layer. Furthermore, the inability to drive the majority of neurons in superficial layer 2/3 of mouse V1 with grating stimuli indicates that there may be fundamental differences in the role of V1 between rodents and other mammals.

## Introduction

The mouse is an excellent model system for taking advantage of the power of two-photon imaging of neural activity. In addition to the power genetic manipulations bring for studying cell-type specific circuitry, their brains are easier to image than larger species such as cats and primates. This is due to their smaller size, which makes it easier to control heartbeat and respiratory pulsations, and to the fact that their brains do not scatter as much light, allowing for deeper imaging. Additionally, because the mouse cortex is much thinner than cats and primates, even with the conventional two-photon imaging depth limit of approximately 400 µm, neurons from layers 1 through 4 can be easily accessed in the mouse visual cortex. In cats and primates, however, two-photon imaging is limited to the upper portion of layer 2/3. Although this ability to image more cortical lamina in the mouse can certainly be an advantage, it raises important considerations about how populations of neural activity are analyzed. The different cortical layers have different anatomical and functional characteristics and play different roles in information processing (Douglas and Martin, 1991). Despite this, most imaging studies in mouse V1 pool neuronal responses without regard for recording depth. Even where cortical lamina is taken into consideration, it has been rare to distinguish neurons by depth within a layer.

In addition to these anatomical differences between mice and higher species, one notable functional difference that has been found is the percentage of responsive neurons in the cortex. Studies have consistently reported that around half of the neurons in mouse V1 respond significantly to grating stimuli (Ebina et al., 2014; Kerlin et al., 2010; Bonin et al., 2011; Mrsic-Flogel et al., 2007; Sohya et al., 2007; Smith and Häusser, 2010; Ayzenshtat et al., 2016; Palagina et al., 2017). However, the few studies done in V1 in primates and cats indicate a much higher percentage of responsive neurons to grating stimuli – typically more than 90% of the cells are responding (Nauhaus et al., 2012; Ikezoe et al., 2013; Kara and Boyd, 2009).

We sought to determine how the response properties of neurons in mouse V1 depend on cortical depth. To ensure that we included the entire neuronal population of each region in our analysis, we used the synthetic calcium indicator Oregon Green BAPTA-1 AM (OGB-1). Although genetically encoded calcium indicators (GECIs) have been engineered that have the sensitivity, dynamic range, and signal to noise levels equal to or exceeding the best synthetic calcium indicators (Chen et al., 2013), they are not ideal for determining the properties of a complete neural population. This is because GECIs do not label all of the cells in a region of tissue (unlike OGB-1) and typical virally-mediated methods give sparse layer 4 labeling. Additionally, the baseline fluorescence of the best GECIs can be so low that cells are invisible when not responding, and so the non-responsive population cannot accurately be determined.

We demonstrate that the percentage of responding and selective neurons, and the amplitude and selectivity of neuronal responses, all vary with imaging depth (even within layer 2/3 itself). Furthermore, we show that although in superficial layer 2/3 only around half of the neurons are responding to grating stimuli, deep in layer 2/3 and in layer 4, nearly all the neurons are responsive. These changes in response properties through depth may be due to differences in the distribution of thalamic inputs terminating in V1. Unlike in cats and primates where the thalamic inputs to the middle layers of cortex are constrained within layer 4 (Gilbert and Wiesel, 1979; Friedlander and Martin, 1989), in mice the inputs ramify extensively into layer 2/3 (Kondo and Ohki, 2016; Sun et al., 2016; Ji et al., 2015; Nakamura et al., 2007). The distribution of terminals shows a gradient of increasing density with depth through layer 2/3 and into layer 4, qualitatively matching the gradient of responsiveness we see.

## Results

We imaged neural activity over depth planes ranging from 150 µm to 350 µm below the surface in 50 µm increments. The density of cells was similar across depth planes except for an increase in the deepest plane (Figure 1, Table 1). Based on the cell count and on the delineation of layers in the literature (Kondo and Ohki, 2016; Sun et al., 2016; Durand et al., 2016; Ji et al., 2015), the three superficial depths (150 µm, 200 µm, and 250 µm) are within layer 2/3, the 350 µm depth is within layer 4, and the 300 µm depth is near the border of layers 3 and 4 and so is difficult to classify. Across all depth planes we found that many cells respond robustly to drifting grating stimuli and they are often sharply tuned for stimulus orientation and direction (Figure 1). However, there was a dramatic increase in the number of cells responding with each increasing depth plane (Figures 1-2). The average percentage of responsive cells in superficial layer 2/3 was approximately 50%, similar to the degree of responsiveness that has been reported in the literature. Deeper in layer 2/3 this rose to ~80%, and in layer 4 around 90% of neurons were responding (*R*^2^ = 0.70, *n* = 35 imaged regions, *P* < 10^−9^, linear regression; Figure 3A, Table 1). This increase in responsiveness with depth was observed in every animal we tested (Figures 2-3A, Table 1; all *R*^2^ > 0.8, *n* = 5 regions per animal, *P* < 0.05).

**Figure 1.**
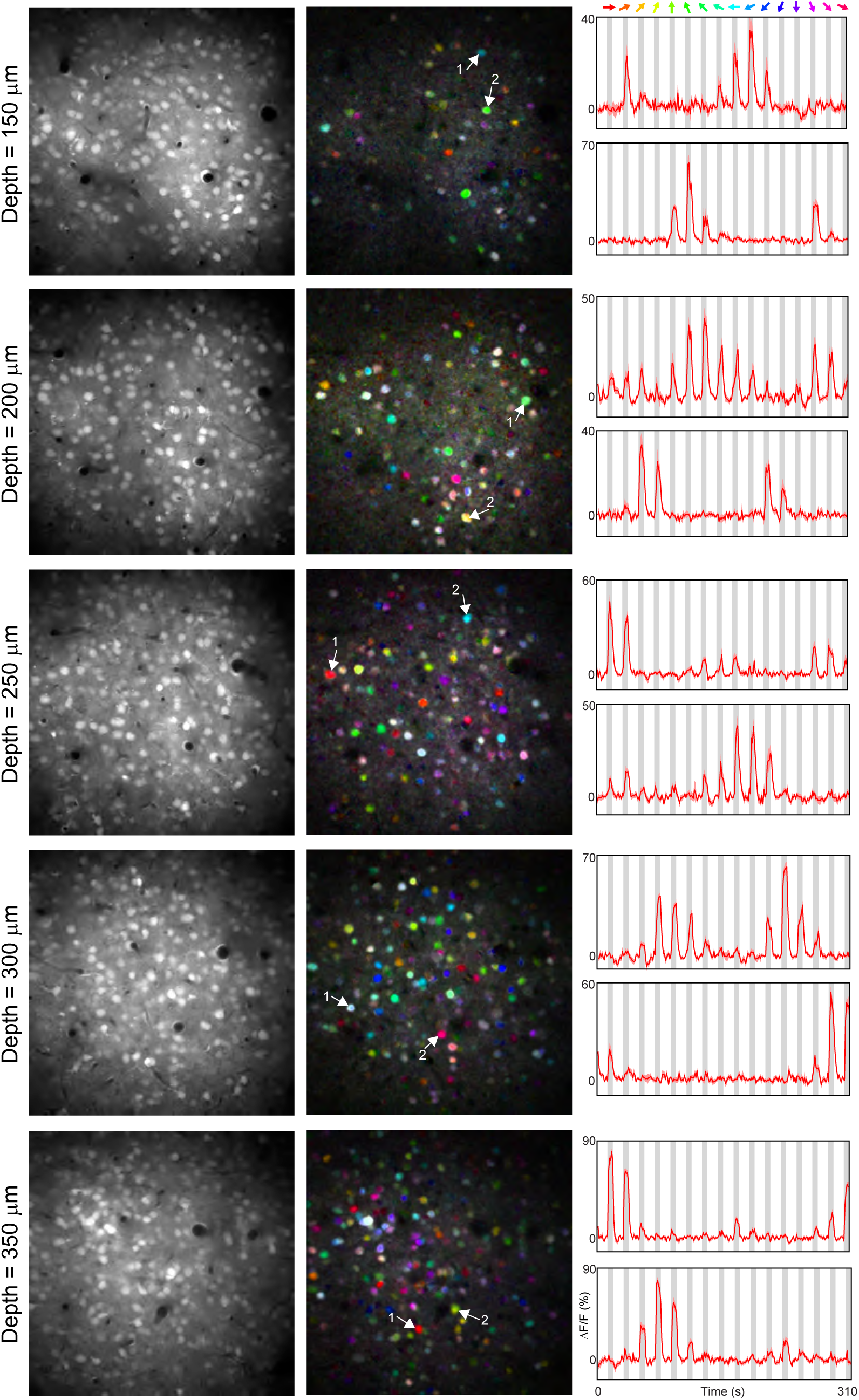
Increased responsiveness with depth in mouse V1. Left column. Anatomical images of five different depth planes from one mouse. Center column. Pixel-based map of stimulus direction. The hue gives the preferred direction, the saturation gives the selectivity, and the brightness gives the response strength. Right column. Times courses of responses from two example cells in each depth plane. The upper traces of each plane correspond to the cells marked #1 in the middle column.

**Table 1.** Summary of data. For the primary parameters of interest, data for individual mice as well as population average data are given for each depth plane. For the bottom two rows, see Methods for a description. The two right columns give the R2 and p values for linear regression on the data. # cells, % Responding, and % Selective data are one value per imaged depth plane. Other data are averages across cells within a depth plane.

| | | 150 μm | 200 μm | 250 μm | 300 μm | 350 μm | R<sup>2</sup> | P |
| --- | --- | --- | --- | --- | --- | --- | --- | --- |
| # Cells | Population average | 181 ± 49 | 176 ± 36 | 179 ± 31 | 186 ± 42 | 237 ± 68 | | |
| % Responding | 160411 | 27 | 55 | 77 | 84 | 80 | 0.81 | 0.0386 |
|  | 160414 | 56 | 66 | 83 | 86 | 93 | 0.94 | 0.0059 |
|  | 160421 | 52 | 72 | 92 | 93 | 94 | 0.82 | 0.0339 |
|  | 160428 | 30 | 55 | 74 | 76 | 79 | 0.83 | 0.0311 |
|  | 160502 | 38 | 58 | 67 | 85 | 91 | 0.97 | 0.0020 |
|  | 160913 | 59 | 77 | 83 | 88 | 95 | 0.92 | 0.0106 |
|  | 160916 | 60 | 73 | 91 | 91 | 92 | 0.81 | 0.0371 |
| | Population average | 46 ± 14 | 65 ± 9 | 81 ± 9 | 86 ± 6 | 89 ± 7 | 0.70 | 4.88E-10 |
| $\Delta F/F$ (%) | 160411 | 12.4 | 11.8 | 13.4 | 19.8 | 22.2 | 0.15 | 0.00E+00 |
|  | 160414 | 13.6 | 14.3 | 16.6 | 22.2 | 25.2 | 0.14 | 0.00E+00 |
|  | 160421 | 15.1 | 19.1 | 19.1 | 23.6 | 31.6 | 0.20 | 0.00E+00 |
|  | 160428 | 10.7 | 13.4 | 12.7 | 18.3 | 20.3 | 0.13 | 0.00E+00 |
|  | 160502 | 12.3 | 13.1 | 13.5 | 18.7 | 26.8 | 0.25 | 0.00E+00 |
|  | 160913 | 14.5 | 17.2 | 16.9 | 21.6 | 27.8 | 0.12 | 0.00E+00 |
|  | 160916 | 16.3 | 15.5 | 17.8 | 24.3 | 33.2 | 0.21 | 0.00E+00 |
| | Population average | 13.6 ± 1.9 | 14.9 ± 2.6 | 15.7 ± 2.5 | 21.2 ± 2.4 | 26.7 ± 4.7 | 0.16 | 0.00E+00 |
| Orientation Selectivity Index | 160411 | 0.73 | 0.50 | 0.51 | 0.54 | 0.51 | 0.02 | 0.001019 |
|  | 160414 | 0.62 | 0.52 | 0.54 | 0.53 | 0.45 | 0.04 | 5.84E-09 |
|  | 160421 | 0.53 | 0.42 | 0.41 | 0.46 | 0.40 | 0.02 | 1.15E-05 |
|  | 160428 | 0.54 | 0.52 | 0.44 | 0.53 | 0.55 | 0.00 | 0.153479 |
|  | 160502 | 0.50 | 0.45 | 0.41 | 0.48 | 0.45 | 0.00 | 0.959027 |
|  | 160913 | 0.66 | 0.56 | 0.53 | 0.54 | 0.50 | 0.03 | 2.55E-06 |
|  | 160916 | 0.65 | 0.56 | 0.52 | 0.55 | 0.54 | 0.02 | 0.000868 |
| | Population average | 0.60 ± 0.08 | 0.50 ± 0.05 | 0.48 ± 0.06 | 0.51 ± 0.03 | 0.48 ± 0.06 | 0.01 | < 10<sup>-10</sup> |
| Tuning Width (degrees) | 160411 | 28 | 25 | 24 | 28 | 31 | 0.02 | 0.003674 |
|  | 160414 | 21 | 25 | 25 | 28 | 38 | 0.15 | 0 |
|  | 160421 | 23 | 27 | 26 | 29 | 36 | 0.08 | 0 |
|  | 160428 | 22 | 23 | 24 | 23 | 27 | 0.02 | 0.002432 |
|  | 160502 | 21 | 22 | 27 | 29 | 32 | 0.08 | 1.33E-10 |
|  | 160913 | 20 | 22 | 26 | 23 | 26 | 0.02 | 0.00139 |
|  | 160916 | 19 | 23 | 25 | 26 | 27 | 0.05 | 4.73E-09 |
| | Population average | 22 ± 3 | 24 ± 2 | 25 ± 1 | 26 ± 3 | 31 ± 5 | 0.06 | < 10<sup>-10</sup> |
| OMI | Population average | 0.80 ± 0.05 | 0.72 ± 0.06 | 0.69 ± 0.07 | 0.73 ± 0.03 | 0.71 ± 0.05 | | |
| % Selective | Population average | 43 ± 14 | 61 ± 9 | 77 ± 10 | 83 ± 6 | 86 ± 8 | 0.69 | 6.26E-10 |
| % Selective of Responding | Population average | 92 ± 4 | 94 ± 2 | 94 ± 3 | 96 ± 2 | 97 ± 2 | 0.26 | 0.001845 |
| % Responding $p \leq 0.001$ | Population average | 38 ± 13 | 57 ± 9 | 73 ± 11 | 81 ± 8 | 85 ± 8 | | |
| % Responding $\Delta F/F \geq 5\%$ | Population average | 55 ± 22 | 76 ± 14 | 86 ± 8 | 90 ± 5 | 93 ± 6 | | |

**Figure 2.**
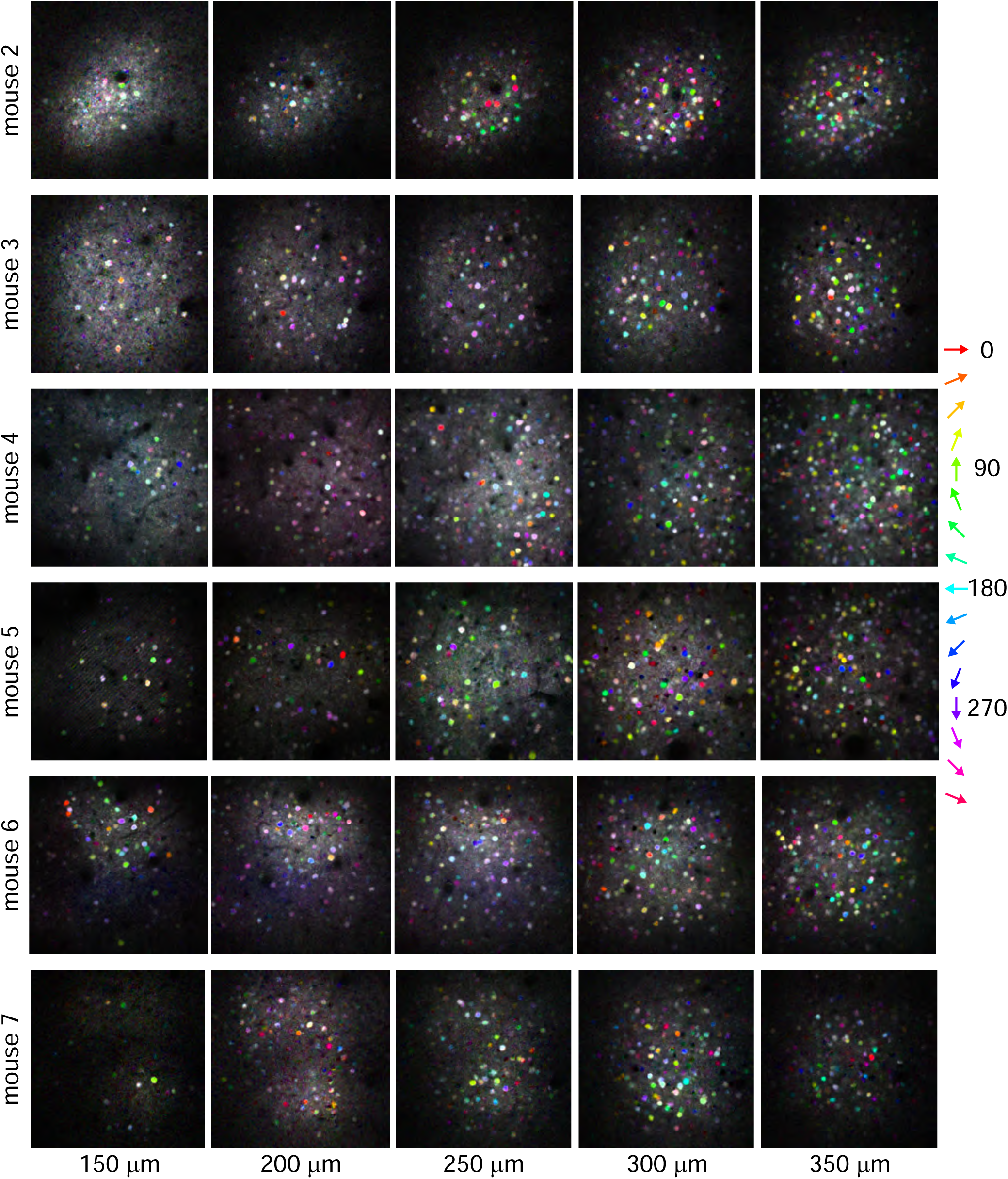
Pixel maps of population. Pixel-based direction maps at each of the five imaging depths from the whole population (excluding the mouse shown in Fig. 1). Color scheme as in Fig. 1.

**Figure 3.**
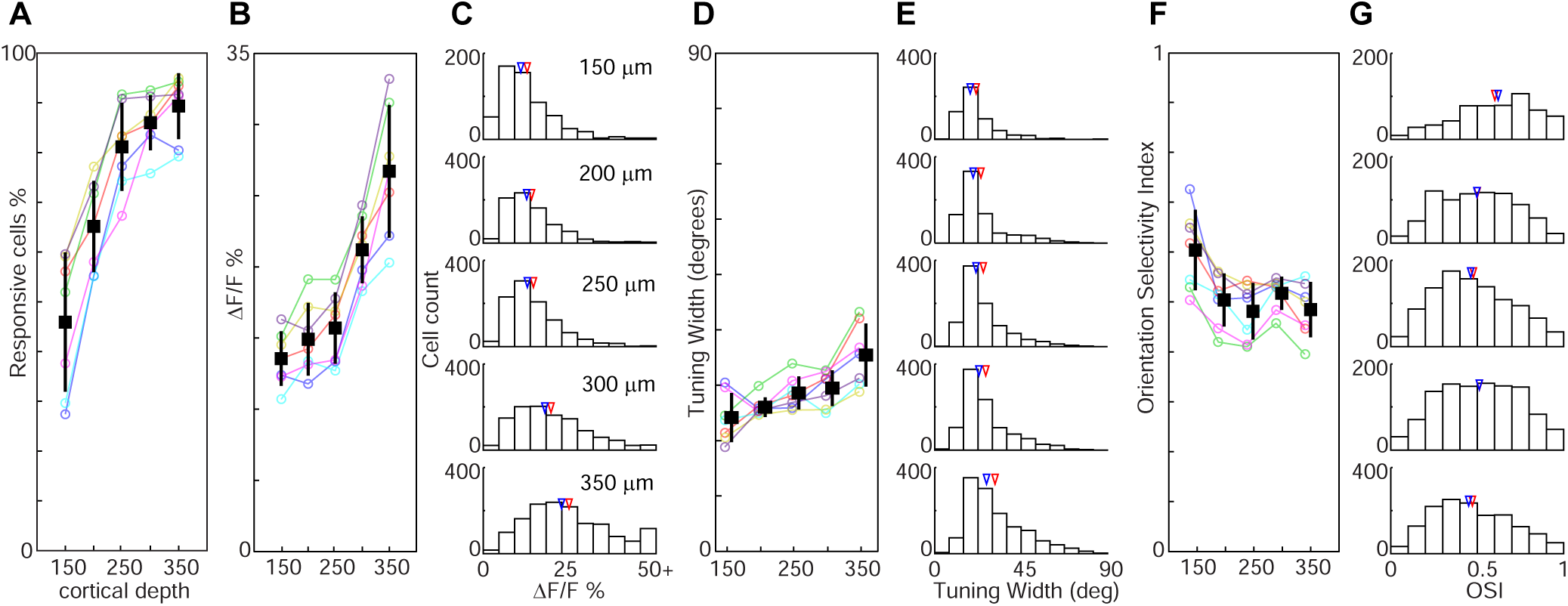
Response properties depend on imaging depth. **A**. The percentage of responding neurons plotted against imaging depth. In this and subsequent panels, colored lines and circles correspond to individual mice and black squares correspond to the population average at each depth. Error bars indicate standard deviation. **B**. Change in response amplitude with depth. **C**. Histogram of the distribution of response amplitude across the population at each depth plane. In this and subsequent panels, blue arrows correspond to (median – check) and red arrows to the (mean – check). **D**. Tuning width versus depth. **E**. Distribution of tuning width values across the population at each depth plane. **F**. Orientation Selectivity Index (OSI) versus depth. **G**. Distribution of OSI values across the population at each depth plane.

We next determined if properties such as the amplitude and selectivity of the responses also showed a depth dependent effect. We compared response amplitudes, computed as the percent change in fluorescence from baseline (% ΔF/F), for the preferred stimulus direction of each neuron. We found that the neuronal response amplitude also steadily increased across depth planes ranging from 13.6 ± 1.9 % (mean ± standard deviation across runs) in superficial layer 2/3 to 26.7 ± 4.7 % in layer 4 (*R*^2^ = 0.16, *n* = 5020 neurons from 7 animals, *P* < 10^−10^; Figure 3B,C, Table 1). This increase in response amplitude with cortical depth was observed in each animal (all *R*^2^ ≥ 0.12, *P* < 10^−10^).

Next, we compared the stimulus selectivity of neurons across depth planes. The percentage of neurons which were selective for stimulus orientation increased with imaging depth, closely following the increase in responsiveness (*R*^2^ = 0.69, *n* = 35 regions, *P* < 10^−9^; Table 1). More than 90% of the responsive neurons in all depth planes were selective for orientation, and there was a slight but significant increase in this value with increasing depth, ranging from 92% in superficial layer 2/3 to 97% in layer 4 (R^2^ = 0.26, *n* = 35 regions, *P* < 0.005; Table 1).

The orientation selectivity of individual cells decreased with increasing depth. The tuning width of responses (half-width at half height; see Methods) showed a continual increase with cortical depth (indicating a reduction in orientation selectivity) from 22 ± 3° in superficial cortical layer 2/3 to 31 ± 5° in layer 4 (*R*^2^ = 0.06, *n* = 4833 neurons, *P* < 10^−10^; Figure 3D,E, Table 1). This trend was observed in each animal (all *R*^2^ ≥ 0.015, *P* < 0.005). The Orientation Selectivity Index (OSI), computed as 1 – Circular Variance (see Methods), also displayed a reduction in selectivity with depth, ranging from 0.60 ± 0.08 in superficial layer 2/3 to 0.48 ± 0.06 in layer 4 (R^2^ = 0.01, *n* = 5020 neurons, *P* < 10^−10^; Figure 3F,G, Table 1). Most of this trend however was due to the drop between the first two planes, and only 5 of the 7 animals showed a significant correlation between OSI and depth (see Table 1). Other imaging studies have emphasized the similarity of orientation tuning between layer 2/3 and layer 4 though they did not compare different depths within each layer (Kondo and Ohki, 2016; Sun et al., 2016). To compare our findings with theirs, we split our data into two groups, categorizing the three superficial depths as layer 2/3 and the two deepest depths as layer 4. The difference in OSI was small, similar to what has been reported (Kondo and Ohki, 2016; Sun et al., 2016) but still highly significant (layer 2/3 vs. layer 4: 0.51 ± 0.23 vs. 0.49 ± 0.23; *P* < 10^−4^, Mann-Whitney test).

Another metric of orientation tuning that has frequently been used in the literature is the difference in response to the preferred and null stimuli divided by their sum: (*R*_pref_ − *R*_null_)/(*R*_pref_ + *R*_null_). We call this the Orientation Modulation Index (OMI) since it gives a measure of the degree to which orientation “modulates” the response (biggest versus smallest response) but it does not fully capture the selectivity of the cells. For instance, as Figure 4A shows, many neurons have the maximum OMI value of 1 but would be considered poorly tuned for orientation by the OSI metric. As long as the response to the null stimulus is essentially zero, the OMI will be 1. But the degree of responsiveness to the intervening orientations can be quite different—when those responses are also close to zero the OSI is very high, but if they are close to the preferred stimulus response, the OSI can be quite low (Figure 4A, data points on right border). Nonetheless, similar to the OSI, the OMI also decreased with increasing depth, with the only notable difference being the consistently higher values of the OMI compared to the OSI (Figure 4B Table 1).

**Figure 4.**
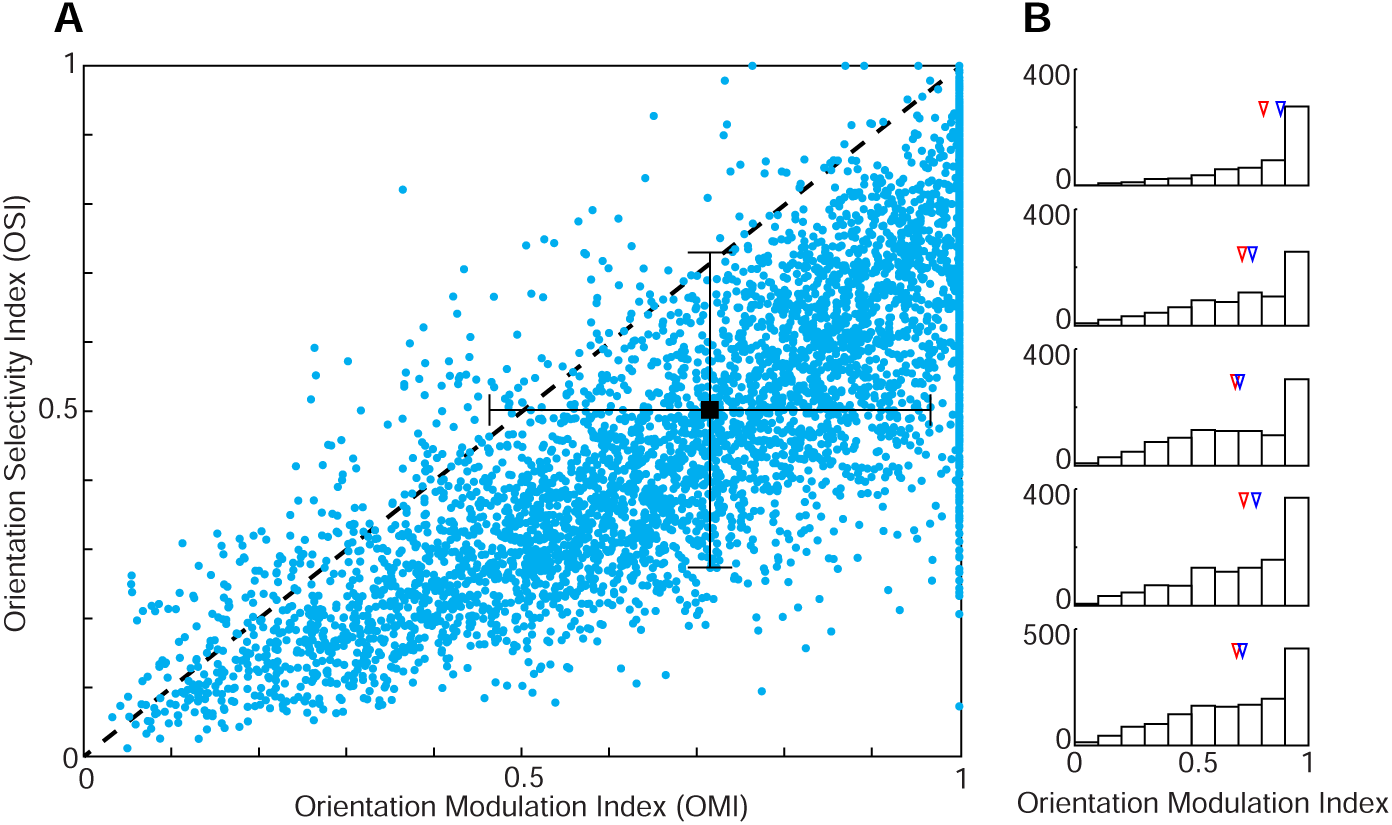
Comparison of OSI and OMI. **A**. Scatterplot showing the OSI and OMI value for all significantly orientation selective neurons. **B**. Distribution of OMI values across the population at each depth plane as for other parameters in Fig. 2.

## Discussion

We have demonstrated that neuronal responses in mouse V1 strongly depend on cortical depth. The percentages of responsive and selective cells and the response amplitude increase with imaging depth from upper layer 2/3 into layer 4, while orientation selectivity decreases over this span. It is important to note that these differences are not simply between layer 2/3 and layer 4 but exist at different depths within layer 2/3 itself. The three superficial depth planes (150 µm, 200 µm, and 250 µm) are within layer 2/3 by any definition, and all of the trends are clearly observed over this span. Additionally, we have shown, for what we believe to be the first time, that it is possible to drive nearly all of the neurons in a region of mouse visual cortex with drifting grating visual stimuli.

### Comparison with previous findings on depth dependence of response amplitude and stimulus selectivity

Prior studies comparing response properties between layer 2/3 and layer 4 of mouse V1 have typically found few differences. While we are not aware of any studies comparing the percentages of responsive and selective neurons, there have been studies comparing response amplitude and selectivity across depth. One electrophysiological study found that in awake mice, evoked firing rates were higher in layer 4 than in layer 2/3 (Dadarlat and Stryker, 2017). However, this same group and others have shown no significant difference in anesthetized mice (Niell and Stryker, 2008; Durand et al., 2016; but see Van den Bergh et al., 2010). In contrast, we find in anesthetized mice a dramatic increase in stimulus-evoked firing rates from layer 2/3 into layer 4. Perhaps differences in anesthesia, neuronal population size, or methodology between these studies and ours account for the different findings.

Orientation selectivity has also been shown to be similar between layer 2/3 and layer 4 in both electrophysiological and imaging studies (Niell and Stryker, 2008; Ma et al., 2010; Van den Bergh et al., 2010; Durand et al., 2016; Kondo and Ohki, 2016; Sun et al., 2016), although one study showed narrower tuning widths in superficial layer 2/3 compared to deep layer 2/3 and layer 4 (Niell and Stryker, 2008). Although not significant, there was a tendency across these studies for greater selectivity more superficially, similar to what we have found. It is also important to note that the reduction in orientation selectivity with depth that we see is a relatively weak effect compared to the increase in the number of cells responding and the response amplitude. Therefore, small differences in other studies might have been overlooked.

### Implications of our results for cortical coding

We believe this is the first time it has been shown that nearly all of the neurons in mouse V1 can be driven by a drifting grating, similar to cats and primates. However, this was only true deeper in the cortex. Thus, a puzzle remains as to why nearly half of the neurons in superficial layer 2/3 are non-responsive. Theories of sparse coding have proposed that neurons in higher areas have more sparse responses than neurons in lower areas because they become selective for increasingly complex stimulus features (Barlow, 1972; Olshausen and Field, 2004). Because mice have simpler visual systems than larger species like cats and primates, functions typically performed by higher areas in those species may be delegated to V1 in the mouse (Niell and Stryker, 2010; Laramée and Boire, 2015; Gavornik and Bear, 2014). For instance, neurons in mouse V1 have been shown to be selective to the pattern motion of a plaid stimulus and not just the motion of the individual components (Palagina et al., 2017; Muir et al., 2015) – a property generally associated with higher visual areas in cats and primates (Movshon et al., 1985; Gizzi et al., 1990; Albright and Stoner, 1995). So perhaps the unresponsive neurons in superficial layer 2/3 would respond to more complex stimuli that would drive neurons in higher areas of other species. Indeed, there are numerous reports in the literature of neurons in mouse V1 whose response selectivity would make them unresponsive to the drifting grating stimuli we used here. For example, cells have been found in mouse V1 that are unresponsive to single gratings but do respond to two overlapping gratings (Palagina et al., 2017; Juavinett and Callaway, 2015). It has also been reported that a contrast-noise stimulus can drive neurons that were unresponsive to gratings (Gandhi et al., 2008; Niell and Stryker, 2008). Additionally, inputs to V1 from other sensory domains, which are essentially non-existent in species like cat and primate, are prevalent in the mouse (Meredith and Lomber, 2017) and so some of the silent neurons may be selective for multisensory inputs. Similarly, locomotion has been shown to dramatically increase firing rates of neurons in V1 (Niell and Stryker, 2010), although whether this also increases the percentage of responsive neurons does not seem to have been determined. But it is reasonable that the increase in response amplitude across the population would lead to more neurons appearing significantly responsive. And finally, neurons have also been found in V1 that only respond to stimuli in the ultra-violet range but not in the visible spectrum (Tan et al., 2015) which again may account for some of the silent neurons. It is important to note however, that regardless of what stimuli might activate these neurons, there is a clear depth dependence in that these alternative stimuli are not necessary for activating the neurons deeper in layer 2/3 and layer 4.

### Circuits for depth dependence of response properties

It may seem surprising that over distances as small as 50 µm there can be such dramatic differences in response properties, particularly when the cells are still in the same layer. One possible source of these differences is the variability in the distribution of thalamic inputs in the upper part of mouse V1. Unlike in other species (Gilbert and Wiesel, 1979; Friedlander and Martin, 1989), the major thalamic input pathway targeting the granular layer in mouse V1 is not constrained to within layer 4 (Kondo and Ohki, 2016; Sun et al., 2016; Ji et al., 2015). Rather, the thalamic inputs spread up into layer 2/3, innervating the deeper part of layer 2/3 quite strongly and becoming sparser more superficially. Thus, as the number of thalamic inputs increases with depth, the feed-forward neuronal drive could also be increasing, leading to a higher percentage of responding neurons and a greater response amplitude. This difference in thalamic inputs could also potentially explain the weakening of orientation selectivity with increasing depth. It has been shown that thalamic inputs are less sharply tuned than neuronal spiking outputs in mouse V1 (Kondo and Ohki, 2016; Sun et al., 2016). Therefore if the broadly tuned thalamic inputs contribute more to the responses of the deeper neurons than the superficial neurons, this could lead to decreased selectivity in the outputs of the deeper neurons. However, as we have noted, the depth-dependence of orientation selectivity is much less dramatic than the effect on responsiveness and response amplitude. Therefore, if the thalamic drive is the explanation for our results, it would appear that there are cortical dynamics – such as spike threshold nonlinearities or local circuitry – that are offsetting the differences created by the thalamic input distribution.

## Methods

Animals and surgery. All surgical and experimental procedures were approved by the Institutional Animal Care and Use Committee at Medical University of South Carolina. C57Bl/6J mice (*n* = 7 male, postnatal day 90-111) were initially anaesthetized with a bolus infusion of fentanyl citrate (0.04–0.05 mg kg^−1^), midazolam (4–5 mg kg^−1^), and dexmedetomidine (0.20–0.25 mg kg^−1^). During two-photon imaging, continuous intraperitoneal infusion with a lower concentration mixture (fentanyl citrate: 0.02–0.03 mg kg^−1^ h^−1^, midazolam: 1.50–2.50 mg kg^−1^ h^−1^, and dexmedetomidine: 0.10–0.25 mg kg^−1^ h^−1^) was administered using a catheter connected to a syringe pump. Craniotomies (2–3 mm square) were opened over the primary visual cortex. A pipette containing a solution with Oregon Green 488 Bapta-1 AM (OGB-1 AM) and a red dye (Alexa 633 or Alexa 594) for visualization was inserted into the craniotomy and the dye was injected with air pressure puffs. The dye loading procedure has been described in detail (O’Herron et al., 2012). After waiting one hour, the dura was removed and the craniotomies were sealed with agarose (1.5–2% dissolved in artificial cerebrospinal fluid) and a 5-mm glass coverslip.

Fluorescence was monitored with a custom-built microscope (Prairie Technologies) coupled with a Mai Tai (Newport Spectra-Physics) mode-locked Ti:sapphire laser (810 nm or 920 nm) with DeepSee dispersion compensation. Excitation light was focused by a 40× (NA 0.8, Olympus) water immersion objective. Full frame imaging of approximately 300 µm square windows was obtained at approximately 0.8 Hz.

Drifting square-wave grating stimuli were presented on a 17-inch LCD monitor. The gratings were presented at 100% contrast, 30 cd m^−2^ mean luminance, 1.5 Hz temporal frequency, and 0.033-0.063 cycles/degree. These stimuli were presented at 16 directions of motion in 22.5° steps (except 1 of 35 runs which used 8 directions in 45 degree steps) for 6.5 seconds with 13 seconds of blank before each stimulus. Each condition was repeated at least 8 times except for two runs with only 5 repetitions.

Images were analyzed in Matlab (Mathworks) and ImageJ (National Institutes of Health). Data with significant movements (several µm) in XY or Z were excluded. Data with small drift movements were realigned by maximizing the correlation between frames. Cell masks were automatically created based on morphological features and then subsequently refined by hand. When using a fully automated cell finding algorithm based on morphology, the overall results showed the same trends. Astrocytes were removed from the data based on morphological criteria (Runyan et al., 2010; Gandhi et al., 2008) and in two animals we verified this method by labeling astrocytes with Sulfrhodamine 101. Fluorescence time courses for each cell were computed by averaging over the pixels in each mask. The time courses were corrected for neuropil contamination similar to Kerlin et al. (Kerlin et al., 2010). First out of focus neuropil contamination was estimated from the fluorescence in small vessels (<15 µm). The fluorescence from hand-drawn vessel masks was divided by the fluorescence of the surrounding neuropil to obtain an estimate of the fraction *C* of the response that is attributable to out of focus contamination. Then the fluorescence time course for soma masks were corrected by subtracting this fraction of the surrounding neuropil fluorescence. So:

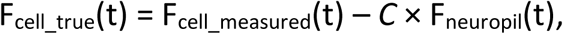

where t is time and F is fluorescence. Values for *C* were between 0.35 and 0.72 (median = 0.56). Neuropil masks were created by expanding a spherical shell 15 µm beyond the soma masks. The inner 3 µm were excluded as a buffer zone around each neuron. Pixels were also excluded from the neuropil masks if they belonged to other soma masks and their 3 µm shells, or were hand drawn as belonging to blood vessels, non-neuronal cell bodies (such as astrocytes), or neuronal somas that were too out of focus to be included in the population. The radius of the neuropil mask was expanded if necessary until the neuropil area was greater than 10 times the soma area. The median radius of the neuropil masks was 14 µm and the range was 12-32 µm.

The time courses for each neuron were then normalized by a sliding baseline of the mean fluorescence of each blank interval (ΔF/F). Responsive neurons were defined by ANOVA across baseline and 16 directions over multiple trials (*P* < 0.01). Because different studies have used different criteria for responsiveness, we also analyzed the data with *P* set to 0.001 and where responsiveness was defined as average ΔF/F > 5% for at least one stimulus direction. The data showed the same depth dependent trends in both cases (see Table 1). Selective neurons were defined by ANOVA across 16 directions over multiple trials (*P* < 0.01). Tuning width was determined by first averaging the responses to the two directions for each orientation. The data were then fit with a least-squares method to a Von Mises function:

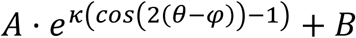

where *θ* is the orientation values, A corresponds to the peak amplitude, *φ* to the preferred orientation, *κ* is a width parameter, and *B* reflects the baseline response (Swindale, 1998). The responses of each neuron were fit 128 times and we divided the full width at the half height of the peak by two to get the half width measure. The mean half width across all 128 fits was used as the tuning width value. The Orientation Selectivity Index (OSI) was defined as:

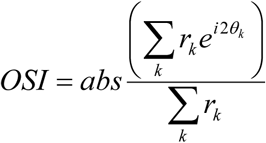

where *θ_k_* is the orientation of each stimulus and *r_k_* is the mean response across trials to that stimulus (Swindale, 1998; Ringach et al., 2002). Note that OSI = 1 – circular variance. The Orientation Modulation Index (OMI) was defined as (*R_pref_* ™ *R_ortho_*)⁄(*R_pref_* + *R_ortho_*), where *R_pref_* is the response to the stimulus direction that evoked the strongest response and *R_ortho_* is the average of the responses to the two orthogonal stimuli. Negative response values (below the baseline level) were set to zero in the computation of OSI and OMI.

All visually responsive neurons were included in the analyses of response amplitude and selectivity except for the case of tuning width where additional criteria had to be met for inclusion in the population. Neurons had to be responsive and selective according to the criteria above (ANOVA) and the preferred stimulus response had to be greater than 3 times the standard error of the mean of the baseline response. Furthermore, if the fit did not converge then the neurons were excluded. We also analyzed the data by including the cells that did not pass the additional criteria by setting their tuning widths to 90 degrees. Although the mean values increased, the trends were the same.

